# An AI/ML-modeled structural atlas of the human protein interactome with functional and cancer-focused networks

**DOI:** 10.1101/2025.07.03.663068

**Authors:** Alexander M. Ille, Christopher Markosian, Stephen K. Burley, Renata Pasqualini, Wadih Arap

## Abstract

In humans, protein-protein interactions mediate numerous biological processes and are central to both normal physiology and disease. While extensive research efforts have aimed to characterize the human protein interactome, atom-scale structural coverage is limited and remains challenging to resolve through experimental methodology alone. Boltz-2, a recent artificial intelligence/machine learning (AI/ML)-based model capable of interaction structure prediction, may serve this experimentally constrained objective. Here, we present *de novo* computed models of binary human protein interaction structures predicted using Boltz-2 based on biochemically determined interaction data sourced from the IntAct database. We assessed the predicted interaction structures through different confidence metrics, examined annotated protein domains with putative interaction involvement, and uncovered interaction networks within the context of biological processes and cancer, highlighting extensive interaction involvement of E3 ubiquitin-protein ligase Mdm2 and p53, among other proteins. This work demonstrates the utility of Boltz-2 for structural modeling of the human protein interactome while also providing novel functional and disease contextualization, holding broad significance for biomedical research at large.

**GRAPHICAL ABSTRACT:** 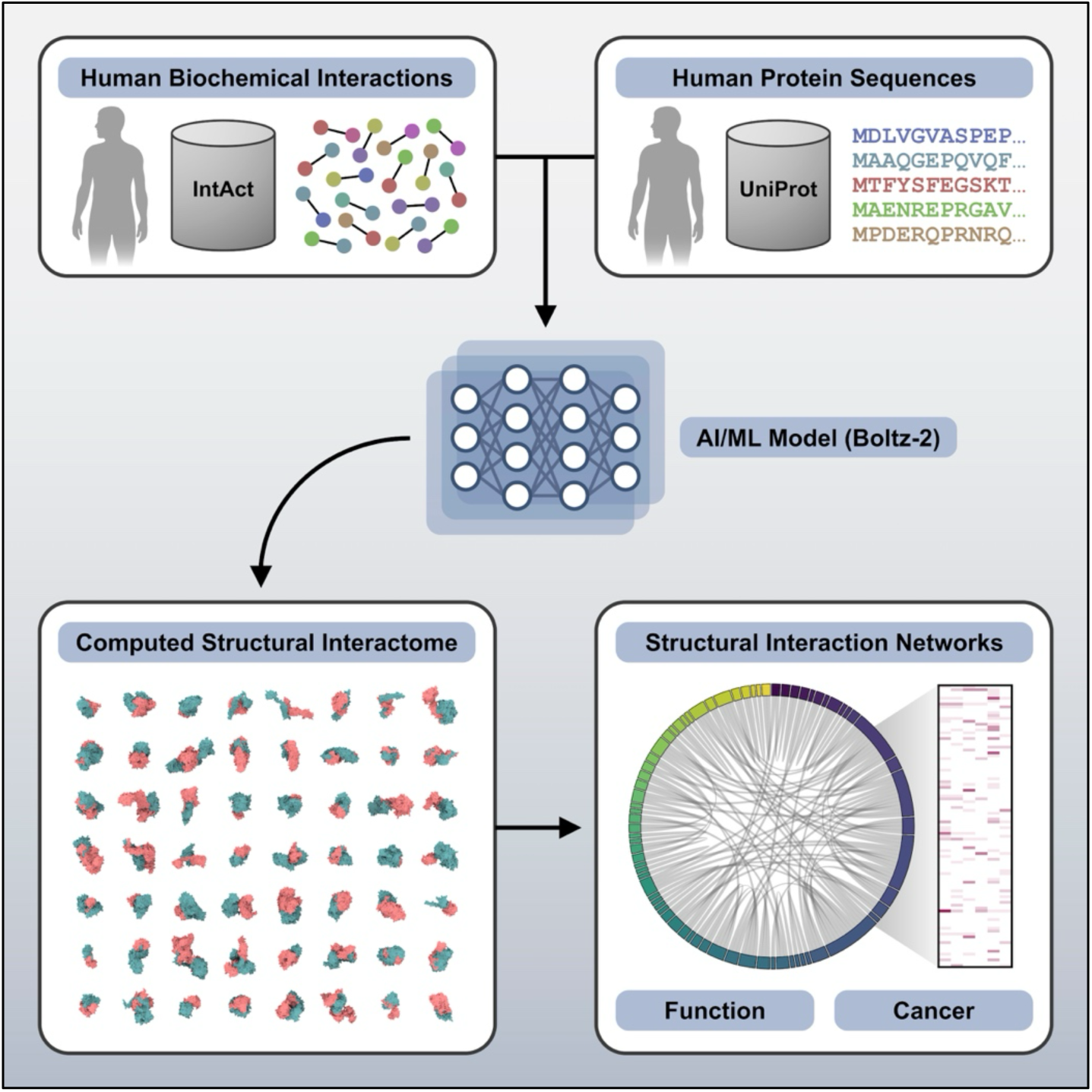

## INTRODUCTION

Interactions between biological macromolecules are fundamental to life, mediating numerous processes in both physiologic and pathological contexts. In particular, protein-protein interactions (PPIs) have been extensively studied for the elucidation of numerous biological functions [1]. Moreover, the identification and characterization of PPIs have long been critical for biomedical research and translational drug development [2, 3]. For example, our group has used *in vivo* phage display [4, 5], an unbiased peptide library screening approach, to uncover a number of functionally actionable interactions [6–13]. PPIs are also commonly probed by using biochemical assays (*e.g.* pull-down, co-immunoprecipitation, mass spectrometry, fluorescence-based techniques) and structural biology approaches (*e.g.* X-ray crystallography, cryo-EM, NMR). Multiple large-scale initiatives have focused on systematic identification and annotation of PPIs based on various forms of experimental evidence [14–19]. These efforts have produced data resources, which provide comprehensive characterization of the ‘interactomes’ of many species including human. In parallel, databases dedicated to sequence annotation and protein structure determination, namely UniProt [20] and the Protein Data Bank (PDB) [21], respectively, have also been critical to systematic profiling of PPIs.

In recent years, metagenomic sequence data and three-dimensional (3D) structure data from open-access repositories [20–23] have enabled development of artificial intelligence/machine learning (AI/ML)-based approaches for protein structure prediction [24–27], including for prediction of structures of biomolecular interactions [26, 28, 29]. AI/ML models trained to leverage the relationship between sequence and 3D structure [30, 31], including those using multiple sequence alignments (MSAs) [24, 26, 28], have exhibited unprecedented accuracy [32, 33] for prediction of both individual structures and those of multiprotein complexes from amino acid sequence alone. AlphaFold-Multimer [29], an initial approach for interaction structure prediction, motivated previous work aimed toward structurally resolving the human interaction network [34]. Other work in this domain has also involved predicted interaction structure-based screening of PPIs [35], large-scale prediction of homo-oligomeric protein assemblies [36], and classifier-guided prediction of human genome maintenance protein interaction structures [37]. Since then, other AI/ML models, including AlphaFold3 [28], RoseTTAFold2-PPI [38], and Boltz-2 [39], have been developed and exhibit substantially improved performance in predicting the structures of protein complexes.

The Boltz-2 model, developed by Saro Passaro, Gabriele Corso, Jeremy Wohlwend et al. [39], has drawn particular interest for its multimodal structure prediction capabilities across a range of biomolecular configurations, including individual chains, multi-chain interaction complexes, and protein-small molecule ligand interactions with corresponding binding affinities. Moreover, Boltz-2 performs robustly in terms of predicting structures of protein-protein, protein-DNA, and protein-RNA complexes, as evaluated against other AI/ML models (including but not limited to AlphaFold3) on a range of complexes freely available from the PDB (following the training cut-off date) [39]. Herein, we leverage the protein-protein structure prediction capabilities of Boltz-2 for large-scale system-level modeling of *n* = 1,394 binary human protein interaction structures, based on biochemically determined PPIs sourced from the IntAct database. The human protein interactome was the focus of the current study due to direct relevance in biomedical research. While other larger-scale human interactome prediction studies have been conducted [34, 38], the current work focuses strictly on interactions supported by biochemical evidence for direct physical contact, thereby providing *a priori* confidence that the predicted structures correspond to genuine binding partners. We assessed predictions using various metrics, considering both overall complex structure and the interaction interface of protein pairs, and found that prediction confidence tended to be greater for smaller complexes while increased MSA depth tended to improve prediction confidence. We also analyzed the modeled complexes for the presence interaction interface-proximal Pfam domains, revealing predicted interaction networks with biological process and oncogenic contextualization from which we highlight extensive interaction involvement of E3 ubiquitin-protein ligase Mdm2 and p53, among other proteins. These findings demonstrate the utility of Boltz-2, highlighting both strengths and limitations, for capturing the broad structural landscape of the human protein interactome and exemplifying opportunities for system-level structural and functional insights with applicability across the biomedical sciences.

## RESULTS

### Large-scale prediction of human protein interaction structures

To construct a representative dataset of the human interactome suitable for structural prediction, we coupled biochemically determined PPIs with their corresponding amino acid sequences. PPIs were sourced from the IntAct database [14] and sequences from UniProt [20], with a binary arrangement of interactor A paired with interactor B **(Figure 1a)**. PPIs were filtered to include only interactions in which both protein interactors were human and designated as having direct physical binding as determined by pull-down assay using purified proteins. Boltz-2 [39] was then used to perform structural predictions of these binary interactions (see Methods). Predictions were performed for protein pairs with individual interactor sequence lengths up to 1,000 residues, providing a dataset consisting of a total of *n* = 1,394 predicted protein interaction structures. For example, the predicted complex structure with the greatest confidence score is presented in **Figure 1b**.

**Figure 1.**
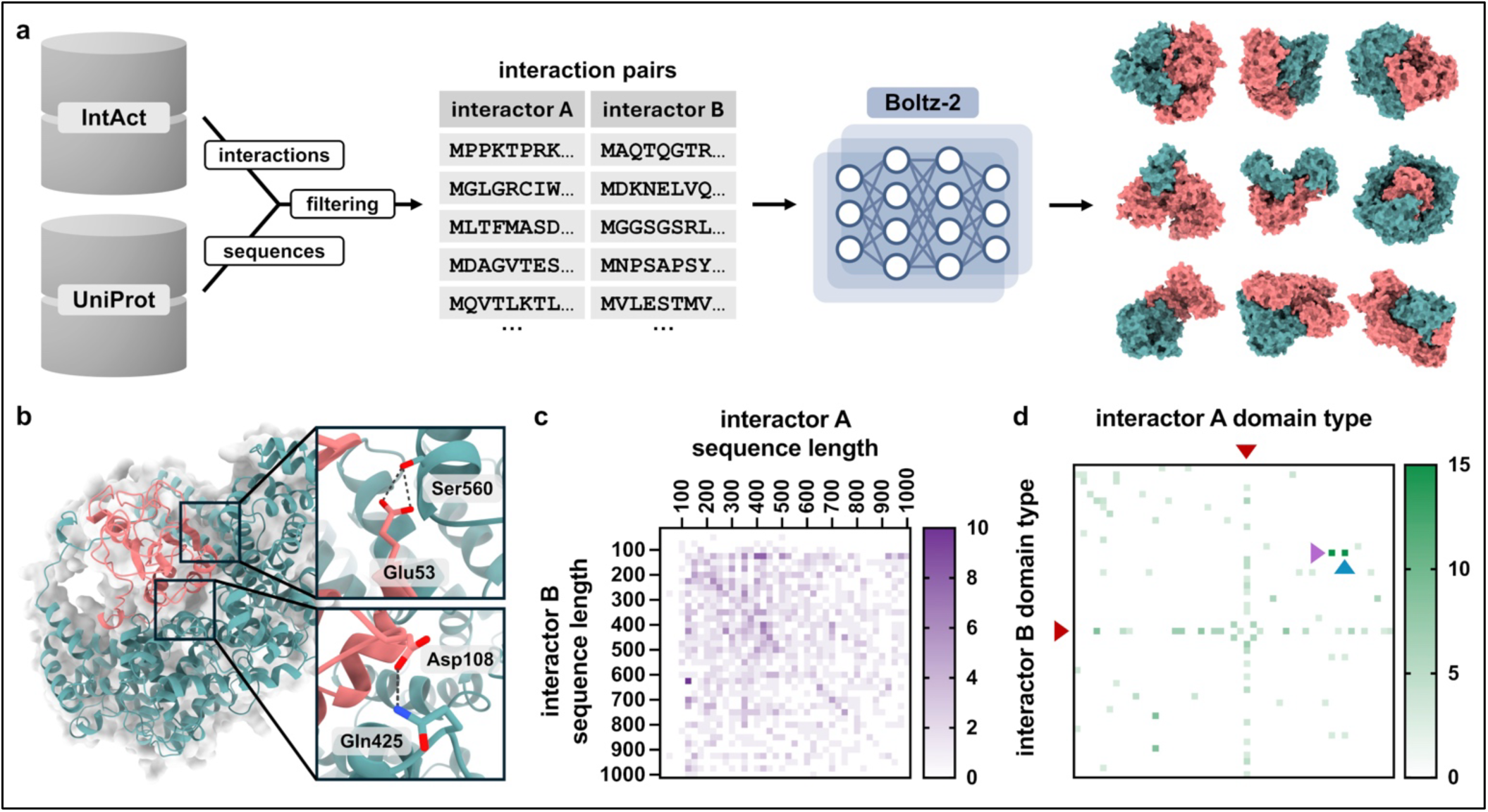
Human protein interactome structure prediction from biochemical interaction and sequence data. **(a)** Overview of protein-protein interaction structure prediction workflow, leveraging human biochemical interaction data and amino acid sequence data from the IntAct [14] and UniProt [20] databases, respectively, for binary complex prediction with Boltz-2. **(b)** Boltz-2 structure interaction prediction with the highest confidence score (0.938) among all interaction structures predicted in the current study. Interactor protein A is depicted in teal, interactor protein B is depicted in red, and dashed lines represent hydrogen bonds. The interaction is that of nuclear cap-binding complex protein subunit 1 and protein subunit 2. Insets show examples of predicted interacting residues between the two proteins. **(c)** Bivariate distribution of sequence lengths of protein interactor pairs, with an upper limit of 1,000 amino acid residues per protein. Data shown as increasing intervals of 25 amino acids, with the scale bar representing number of protein interactor pairs. **(d)** Co-occurrence matrix of Pfam domains across protein interactor pairs. Red arrowheads indicate the most abundant domain, autophagy protein Atg8 ubiquitin like domain (Pfam PF02991), *n* = 215 co-occurrences. Purple arrowhead indicates co-occurrence of the chromatin organization modifier domain (Pfam PF00385) and the zinc finger C3HC4 type RING 2 domain (Pfam PF13923), *n* = 15 co-occurrences, while blue arrowhead indicates co-occurrence of the chromatin organization modifier domain (Pfam PF00385) and the RAWUL domain RING finger- and WD40-associated ubiquitin-like domain (Pfam PF16207), *n* = 15 co-occurrences. Data shown as bins with a minimum of three co-occurrences per bin, with the scale bar representing number of domain co-occurrences.

With the scope of better characterizing the set of predicted interaction structures, we explored sequence-level attributes of the interacting proteins **(Supplementary Table 1)**. Bivariate analysis of the sequence lengths across the protein interactor pairs used for structural prediction revealed a relatively uniform distribution **(Figure 1c)**. This pattern did not indicate any strong bias toward specific sequence length combinations other than most sequence pairs having lengths greater than 100 amino acid residues. For additional biological context, the interactor pair sequences used for structural prediction were queried against the Pfam database [40, 41] to identify co-occurrence of annotated protein domains **(Figure 1d)**. In total, there were 2,391 unique domain co-occurrences, highlighting rich biological variability among the predicted structures. The most abundant co-occurring domains were the chromatin organization modifier domain (Pfam PF00385) with the zinc finger C3HC4 type RING 2 domain (Pfam PF13923) and the chromatin organization modifier domain (Pfam PF00385) with the RAWUL domain RING finger- and WD40-associated ubiquitin-like domain (Pfam PF16207), each present in 15 protein interactor pairs. Notably, the autophagy protein Atg8 ubiquitin like domain (Pfam PF02991) was present in 173 protein interactor pairs. It should be noted that the presence of these domains does not imply interaction involvement in the interactor pairs in which they are found—*i.e.* these domains may or may not be located at the interaction interface, which is addressed with further analysis below.

### Prediction confidence and sequence-dependent effects

To gain insight into the quality of the predicted interaction structures, we used confidence metrics for assessment of overall structure as well as the interaction interface **(Supplementary Table 2)**. Predicted local distance difference test (pLDDT) [24, 42] and predicted template modelling (pTM) [24, 43] scores were used for evaluation of overall structure. In brief, pLDDT scores residue-level confidence while pTM scores confidence in regional topology. Interaction interface-restricted versions of pLDDT and pTM were used for assessment of interaction-specific structural confidence [28, 29, 39]. The Boltz-2 confidence score, which combines overall pLDDT with interface pTM, was also included for evaluation [39] (see Methods). These metrics each exhibited a wide range of values across the dataset of predicted interaction structures **(Figure 2a)**. The distribution of these values suggests greater residue-level confidence (as indicated by pLDDT metrics) and lower confidence in regional topology (as indicated by pTM metrics), while the combined confidence score had a median value of 0.583, indicating that more than half of the structures were predicted with moderate confidence.

**Figure 2.**
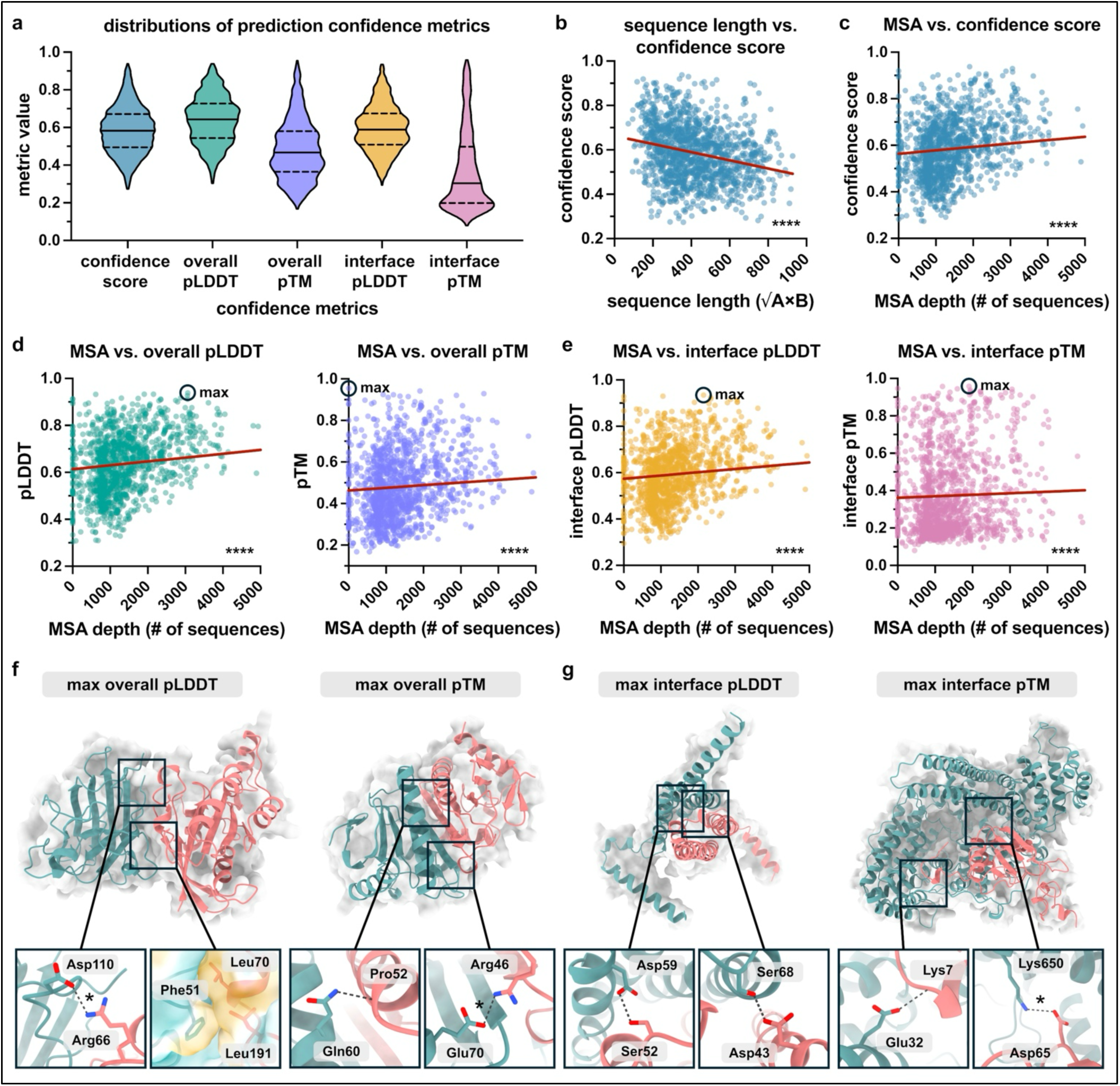
Confidence metrics of predicted interaction structures. **(a)** Prediction confidence metric distributions across all protein interaction structures predicted (*n* = 1,394). All metrics were determined by Boltz-2 on a normalized scale of 0 to 1, enabling direct comparison (see Methods). Solid lines represent medians and dashed lines represent first and third quartiles. **(b)** Relationship between combined sequence length (the geometric mean of interactor sequence A and interactor sequence B) and confidence score of the predicted interaction structures. Spearman correlation *r* = -0.261, \*\*\*\**p* <0.0001. **(c)** Relationship between MSA depth and confidence score of the predicted interaction structures. Spearman correlation *r* = 0.353, \*\*\*\**p* <0.0001. Predicted interaction structures with MSA depth greater than 5000 (*n* = 25 structures) are not graphically depicted here, but were included in statistical analyses for panels c-e. **(d)** Relationship between MSA depth and entire interaction complex (overall) pLDDT and pTM. Spearman correlation *r* = 0.369 for overall pLDDT, Spearman correlation *r* = 0.266 for overall pTM, \*\*\*\**p* <0.0001. **(e)** Relationship between MSA depth and interaction interface-aggregated pLDDT and pTM. Spearman correlation *r* = 0.331 for interface pLDDT, Spearman correlation *r* = 0.139 for interface pTM, \*\*\*\**p* <0.0001. Lines determined by linear regression are depicted in red in panels b-e. **(f-g)** Predicted interaction structures which exhibited the greatest overall (f) and interface (g) metrics. Interactor protein A is depicted in teal and interactor protein B is depicted in red. Dashed lines represent hydrogen bonds, dashed lines with an asterisk represent salt bridges, and yellow surfaces represent hydrophobic regions.

We also considered whether sequence length and MSA depth might influence prediction confidence. The relationship between combined sequence length of interaction structures (defined here as the geometric mean of the two interactor proteins) and the overall confidence score indicates that structures with shorter sequences were predicted somewhat more confidently, though this negative correlation was relatively weak (*r* = -0.261) **(Figure 2b)**. Comparing MSA depth with the overall confidence score revealed a weak-to-moderate positive correlation (*r* = 0.353), suggesting that increased MSA depth may improve prediction confidence **(Figure 2c)**. Further analyses comparing MSA depth with overall **(Figure 2d)** and interface **(Figure 2e)** pLDDT and pTM metrics showed varying degrees of prediction confidence improvement with increased MSA depth, although relatively limited in correlation strength. The interaction network of protein complexes within the first (upper) quartile of the confidence metric **(Figure 1a)**, additionally limited to proteins involved in at least two binary interactions, is depicted in **Figure 3a**, along with examples of corresponding predicted structures **(Figure 3b & c)**.

**Figure 3.**
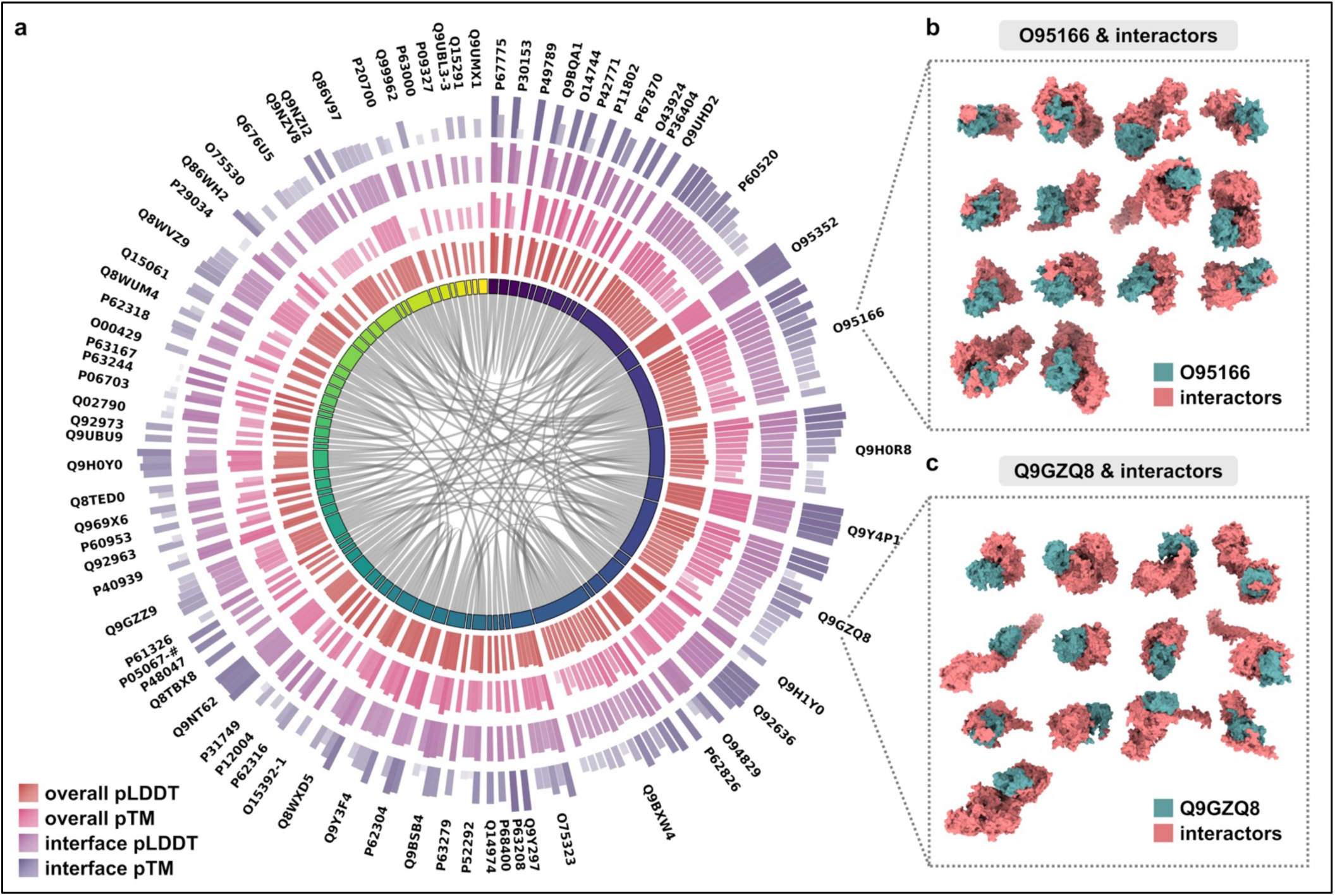
Network of protein interaction structures predicted with high confidence. **(a)** Chord diagram depicting the interaction network between proteins with a minimum of two other interactors where the interaction confidence score is greater than 0.6709 (third quartile, Figure 2a). Individual proteins are plotted along the inner circumference of the circle, with differing colors for each protein. Grey connecting lines (chords) represent interactions between proteins. The four outer layers consisting of scalar bars denote (1) overall pLDDT, (2) overall pTM, (3) interface pLDDT, and (4) interface pTM. For space conservation, P05067-# is used as a short-form for P05067-PRO_0000000092. **(b-c)** Predicted interaction structures for the two proteins with the greatest number of interactors from (a). This includes *n* = 14 binary interactions for γ-aminobutyric acid receptor-associated protein (UniProt O95166) (b) and *n* = 13 binary interactions for microtubule-associated protein 1 light chain 3 β (UniProt Q9GZQ8) (c).

We next performed a post-training evaluation of prediction accuracy using experimentally resolved human protein structures as a reference. Given that the training cutoff for Boltz-2 was June 1, 2023, we searched the PDB [21] for structures deposited after the cutoff date for entries consisting of human binary protein complexes corresponding to those predicted. While only three such structures were identified, PDB IDs 9B4Y, 8X8A [44], and 8T1H [45], they provide a means for experimental comparison. These experimentally determined complex structures were compared to the corresponding interaction structures predicted by Boltz-2 and by AlphaFold3 **(Supplementary Figure 1)**. DockQ [46] scores and interface RMSD were computed between experimentally determined and predicted complex structures, with the predicted structures generated by both models ranging in scores from medium quality (DockQ of 0.49 – 0.80) **(Supplementary Figure 1a)**, acceptable quality (DockQ of 0.23 – 0.49) **(Supplementary Figure 1b)**, and incorrect (DockQ of 0 – 0.23) **(Supplementary Figure 1c)**. Together, these results highlight various strengths and limitations of large-scale protein interaction structure prediction with Boltz-2, including favorable residue-level prediction confidence contrasted by relative uncertainty in regional topology prediction. Furthermore, the availability of additional sequence context through increased MSA depth, though not found to be a requirement for high prediction confidence, may have a moderately positive effect.

### Interaction interface domains, biological process, and oncogenic involvement

The presence and co-occurrence of Pfam domains [40, 41] uncovered during our sequence-level analyses **(Figure 1d)** motivated us to explore the putative interaction involvement of these domains based on the 3D structural complexes predicted with Boltz-2. In order to identify Pfam domains at the interaction interface, we employed a standard < 5 Å distance cutoff [47, 48] between any atom in a given domain present in one interactor protein and any atom in the entirety of the other interactor protein. Under this proximity criteria, a Pfam domain was considered to be putatively involved in a protein-protein interaction. Analysis of the predicted interaction structures revealed that 679 structures (48.7%) contained Pfam domains within 5 Å of the interaction interface compared to 793 structures containing Pfam domains without any proximity restriction **(Supplementary Table 3)**. The prevalence of Pfam domains varied considerably **(Supplementary Figure 2a, Supplementary Table 4)**, with the autophagy protein Atg8 ubiquitin like domain (PF02991) found in the greatest number of interaction structures within 5 Å of the interaction interface. Individual examples of prevalent domains from within the predicted interaction structures are shown in **Supplementary Figure 2b**.

We also analyzed the co-occurrence of Pfam domains that were present within 5 Å of the interaction interface **(Figure 4a-d)**. The autophagy protein Atg8 ubiquitin like domain (Pfam PF02991) had the greatest level of co-occurrence with other domains, aligning with the high prevalence of this domain across interaction structures **(Supplementary Figure 2a)**, while the chromatin organization modifier domain (Pfam PF00385) co-occurred most frequently with the zinc finger C3HC4 type RING 2 domain (Pfam PF13923) and the chromatin organization modifier domain (Pfam PF00385) co-occurred most frequently with the RAWUL domain RING finger- and WD40-associated ubiquitin-like domain (Pfam PF16207). While the structural interaction interface proximity-restricted domain co-occurrences were less prevalent overall, the general pattern of co-occurrence resembled that observed from sequence-level analysis **(Figure 1d)**. Additionally, we sought to infer a broad contextualization of biological function based on the predicted structural interactions paired with interaction interface proximity-restricted domain annotation. As a means for high-level functional categorization, we utilized UniProt biological process keywords—a controlled vocabulary augmented with Gene Ontology (GO) annotation [49, 50] **(Supplementary Table 3)**. Six broad biological process groupings of the domain-based interactions were compiled, including: apoptosis, cell division, differentiation, DNA repair, immunity, and protein transport. These groupings reveal intricate domain-based interaction networks across the predicted structural complexes **(Figure 4e)**. Furthermore, these groupings varied in Pfam domain prevalence and overlap **(Supplementary Figure 3)**, indicating potential structural involvement of specific domains within and across biological processes.

**Figure 4.**
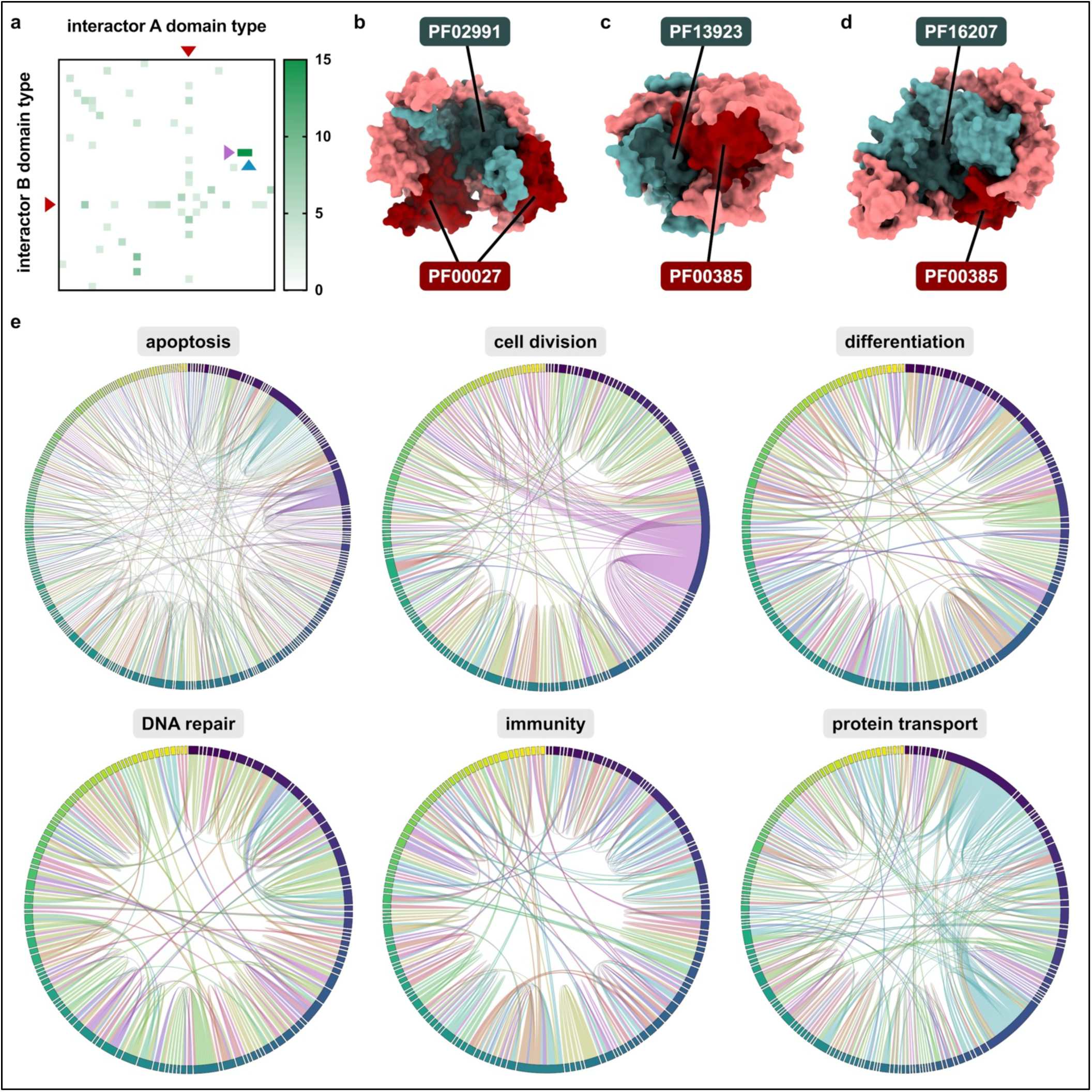
Domain co-occurrence and domain-based biological process interaction networks. **(a)** Co-occurrence matrix of Pfam domains within 5 Å of the interaction interface between protein interactor pairs. Red arrowheads indicate the most abundant domain, PF02991 (*n* = 122 interaction interface co-occurrences), purple arrowhead indicates co-occurrence of domains PF13923 and PF00385 (*n* = 15 interaction interface co-occurrences), and blue arrowhead indicates co-occurrence of domains PF16207 and PF00385 (*n* = 15 interaction interface co-occurrences). Data shown as bins with a minimum of three protein interactor pairs per bin, with the scale bar representing number of protein interactor pairs. **(b-d)** Examples of predicted interaction structures which include co-occurrences of domains PF02991 with PF00027 (b), PF13923 with PF00385 (c), and PF16207 with PF00385 (d). **(e)** Chord diagrams illustrating domain-based interaction networks grouped by UniProt biological process keyword annotation. For each diagram, individual proteins are plotted along the circumference of the circle and connecting lines (chords) represent interactions with other proteins involving domains < 5 Å of the interaction interface. Differing colors along the circle circumference represent unique proteins while differing colors of the chords represent unique domains. UniProt ‘biological process’ keywords used for grouping include KW-0053 (apoptosis), KW-0132 (cell division), KW-0221 (differentiation), KW-0234 (DNA repair), KW-0391 (immunity), and KW-0653 (protein transport), with at least one protein in the interaction pair annotated with the designated keyword required for group inclusion.

Another aspect we considered was domain-based pathological involvement of the predicted protein complexes. We focused on involvement in cancer, specifically oncogenic proteins. Filtering the predicted structures by the UniProt disease keyword “proto-oncogene” (KW-0656), with either one or both interactors containing this annotation, revealed an interaction network consisting of *n* = 129 potentially oncogenic interactions **(Supplementary Table 5, Figure 5a)**. Furthermore, the majority of proteins involved in these interactions have genetic variants, as annotated in the Single Nucleotide Polymorphism Database (dbSNP) [51], with missense mutations involving amino acids at the interaction interface (within 5 Å of the interacting protein). Notably, the protein with the greatest number of interactors within this network was found to be the E3 ubiquitin-protein ligase Mdm2 (UniProt Q00987), and for each of the 12 interactions involving Mdm2, the zinc finger C3HC4 type ‘really interesting gene’ (RING) 3 domain (Pfam PF13920) was present at the interaction interface **(Figure 5b)**. Mdm2 is overexpressed in over 40 cancer types [52] and its oncogenic activity has been characterized to involve inhibition of p53 by ubiquitination via its RING domain [53–57]. This aligns with p53 (UniProt P04637) being among the Mdm2 interactors identified within the oncogenic network and the presence of the RING domain at the structural interface of this interaction **(Figure 5b)**. There is also evidence for cancer involvement among several of the other Mdm2 interactions found in the oncogenic network, including with nucleolin (UniProt P19338) [58, 59], nucleophosmin (UniProt P06748) [60, 61], β-arrestin 2 (UniProt P32121) [62, 63], elongation factor 1α (UniProt P68104) [64, 65], as well as ribosomal proteins S7 (UniProt P62081), S27 (UniProt P42677), and S27L (UniProt Q71UM5) [66–69]. The multiple Mdm2 interactions identified in the oncogenic network reinforce the notion and accumulating evidence that the role of Mdm2 in cancer extends beyond its interaction with p53 [56, 57].

**Figure 5.**
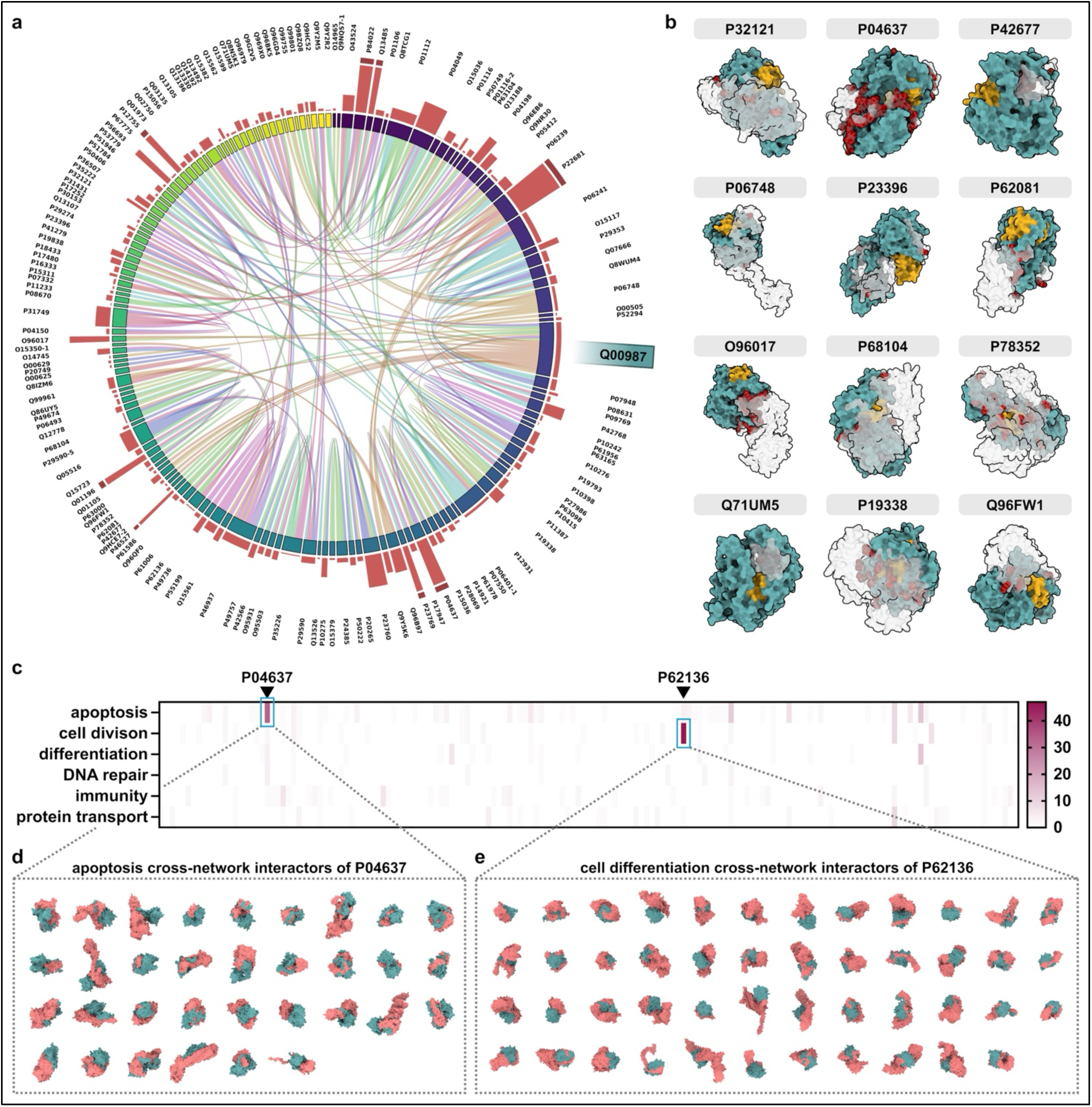
Oncogenic protein interaction network inferred from domain-based interactions. **(a)** Chord diagram depicting an oncology-focused interaction network based on domains proximal to the interaction interface. As in Figure 4e, the circumference of the circle contains individual proteins (differing by color) with connecting lines (chords) representing interactions between proteins which involve domains within 5 Å of the interaction interface. Interaction pairs were included on the basis of at least one protein being annotated with the UniProt disease keyword proto-oncogene (KW-0656). The red outer layer consisting of scalar bars indicates the average number of documented missense mutations located at the various interaction interfaces of each protein, ranging from 0 to 100 mutations with averages greater than 100 denoted by dark red bars. E3 ubiquitin-protein ligase Mdm2 (UniProt Q00987), the protein with the greatest number of interactors within the oncogenic network, is outlined in a teal box. **(b)** Structures of Mdm2 (teal) and its various interacting proteins (transparent) from within the oncogenic network. The zinc finger C3HC4 type RING 3 domain (Pfam PF13920) of Mdm2 is depicted in yellow and missense mutations of the interacting proteins present within 5 Å of the interaction interface are depicted in red. **(c)** Cross-network matrix of proteins mutually present within the oncogenic network and the biological process networks (apoptosis, cell division, differentiation, DNA repair, immunity, and protein transport), with the scale bar representing number of interactions involving each mutually present protein within the respective biological process networks. The mutually present proteins p53 (UniProt P04637) and PPP1CA (UniProt P62136) with the greatest number of interactions in the apoptosis and cell division networks, respectively, are outlined with blue boxes. **(d-e)** Corresponding cross-network interaction structures involving p53 from the apoptosis network (d) and PPP1CA from the cell division network (e) are shown, with p53 and PPP1CA depicted in teal and their interactor proteins depicted in red.

In order to further integrate these data, we performed cross-network analyses between the oncogenic network and the biological process networks **(Figure 5c)**. Proteins mutually present in both the oncogenic network and each respective biological process network with the greatest number of interactions in the latter are presented in **Figure 5d-e** and **Supplementary Figure 4**. The cross-network proteins p53 (UniProt P04637) **(Figure 5d)** and serine/threonine-protein phosphatase 1 catalytic subunit α (PPP1CA) (UniProt P62136) **(Figure 5e)** had the greatest number of interactions in the apoptosis and cell division networks, respectively (*n* = 33 apoptosis network interactions involving p53 and *n* = 47 cell division network interactions involving PPP1CA). Cross-network proteins with the greatest number of interactions in their respective biological process networks, although relatively fewer than found for apoptosis and cell division, included sirtuin-1 (UniProt Q96EB6) for differentiation, p53 (UniProt P04637) for DNA repair, tyrosine-protein kinase Fyn (UniProt P06241) for immunity, and programmed cell death 6 (PDCD6)-interacting protein (UniProt Q8WUM4) for protein transport **(Supplementary Figure 4)**. The cross-network interaction participation of p53 in both apoptosis and DNA repair is particularly consistent with extensive research characterizing the involvement of p53 in these processes [70, 71] while also emphasizing the multi-functional complexity of p53, which remains incompletely understood [72, 73]. Collectively, these structure-based network analyses yield an extensive set of novel data that situate protein-protein interaction profiles within functional and oncogenic contexts, while providing an *in silico* resource for future research into previously unexplored structure-function relationships across broad structural coverage.

## DISCUSSION

This work demonstrates the utility of Boltz-2 for structural modeling of the human protein interactome through large-scale prediction of binary interaction structures based on biochemical data and presents subsequently revealed structure-based interaction networks representing biological function and cancer. Although predictions performed in the current work were derived from an experimentally-constrained subset of human PPIs, the scalability of this approach supports application to larger datasets. Our analyses revealed that over half of the structures were predicted with moderate confidence, and that increased MSA depth was associated with greater prediction confidence, though only moderately. Another trend we observed, albeit weakly correlated, was decreased prediction confidence with increased sequence length, indicating a potential limitation for the prediction of larger protein complexes. Notably, in the post-training evaluation comparing predicted complex structures to experimentally determined structures from the PDB, Boltz-2 and AlphaFold3 performed strongly and struggled on the same cases, *i.e.* similar results were observed for the same proteins. This observation could indicate that certain structural classes or interface types remain inherently challenging to predict. Novel training strategies and architecture refinements may serve to improve predictive accuracy for currently challenging targets. Nevertheless, caution should be exercised when interpreting predicted complex structures generated with either computational tool, and confidence metrics may be useful indicators in this regard.

Our investigation of annotated domains and functional contextualization were also insightful. The finding that 48.7% of the predicted structural complexes contained Pfam domains within 5 Å of the interaction interface highlights the importance of domains in protein-protein interactions [74, 75]. The domain-based interactions inferred in the current work, although predictive, provide a broadened representation of interaction involvement given the substantially increased structural coverage afforded through structural prediction. Furthermore, the elucidation of domain-based interaction networks categorized by biological process provides system-level overviews of interaction intricacy and variability with respect to structure-function relationships. Similarly, prediction-informed network analysis of oncogenic proteins revealed both system-level structural complexity and interaction interface presence of mutation-prone residues corresponding to annotated genetic variants. Despite the inherent complexity, disease-relevant insights may be extracted from this network data, as exemplified by the various Mdm2 interactions with prior evidence for cancer involvement and cross-network analyses.

The relationship between Mdm2 and p53 has been a long-standing subject of research [53, 54, 56, 57, 76–79], initially considered as relatively straightforward—yet continued investigation has revealed multifactorial interplay involving the two proteins for which our understanding remains incomplete [56, 57, 70, 72, 73]. The Mdm2 interactions identified within the oncogenic network **(Figure 5a and b)** exemplify this complexity while in the same vein provide insight in terms of prediction-informed identification and structural characterization of interactor protein candidates. It was also interesting that the RING domain was found at the interface of all 12 Mdm2 interactions, given that this domain mediates the ubiquitination activity of Mdm2 [53–57]. Beyond this, the cross-network analyses pointed to differing interaction profiles within the biological process networks of proteins mutually present in the oncogenic network **(Figure 5c)**, with certain proteins highlighted for extensive interaction involvement **(Figure 5d and e, Supplementary Figure 4)**. These system-level network characterizations are enabled by increased structural coverage, and may serve as a basis for future work aiming to establish novel structure-function relationships and better understand pathologic processes.

Looking ahead, it is reasonable to propose that Boltz-2 could be used to model multi-protein complexes (*i.e.* higher-order assemblies) of the human interactome, going beyond the binary interaction modeling performed in the current study. Boltz-2 is capable of performing predictions of structures with more than two chains, and large-scale modeling has previously been explored for protein complexes with AlphaFold2-based approaches [34, 36, 37]. Moreover, the insights derived from both structural characterization and subsequent network analyses may be informative for guiding targeted therapeutic development. Multiple groups [80–83], including ours [10, 12, 84], have incorporated computational approaches to structurally probe protein interactions relevant to therapeutic development, and we anticipate large-scale interaction structure predictions based on biochemical data as presented herein will similarly play a key role. The current work suggests Boltz-2 is a valuable tool for human interactome protein structure prediction and related work, though results should be interpreted with appropriate consideration of current limitations. Emerging AI/ML-based approaches for modeling the structures of interactions and corresponding system-level analyses are anticipated to aid multiple areas of biomedical research through broadened characterization of complex biomolecular landscapes.

## METHODS

### Curation and processing of interaction and sequence data

The *Homo sapiens* species-specific dataset, which lists binary interaction pairs by UniProt ID, was downloaded from the IntAct database in tabular format [14]. The dataset was filtered as follows: (1) removal of all interaction pairs for which one of the two interacting proteins were non-human; (2) exclusive retention of interaction pairs which contained the “Direct interaction” designation; and (3) exclusive retention of interaction pairs which determined by pull down as the experimental interaction detection method. Corresponding sequences of interaction pairs were obtained by UniProt ID through the UniProt website REST API [20]. Corresponding Pfam domains were obtained through the InterPro Rest API [40]. Twelve interaction pairs were excluded from data analysis. Three interaction pairs were excluded due to their sequences containing the amino acid selenocysteine. Additionally, nine interaction pairs (P54278 with P40692; Q9UBS5 with P46459; Q8N8A2 with P35125-3; Q9UIF7 with P43246; Q13563 with Q13563; Q92622 with Q8NEB9; P11142 with P34932; Q04759 with Q8IVH8; and Q9UNQ0 with P22413) for which Boltz-2 failed to generate predictions despite multiple attempts, were also excluded. With all of the above curation and exclusion, a total of *n* = 1,394 binary interaction pairs was analyzed in the current work.

### Interaction structure prediction and analyses

Structure predictions were performed using Boltz-2 [39] version 2.1.1 in the Google Colab environment. Hardware configurations were set to A100 GPU and High-RAM. The paired interaction protein sequences were used as input for protein complex prediction. Default Boltz-2 settings were used, except for MSA server use set to True, as follows:

diffusion_samples = 1

recycling_steps = 3

sampling_steps = 200

step_scale = 1.638

use_msa_server = True

msa_server_url = https://api.colabfold.com

max_msa_seqs = 8192

subsample_msa = False

msa_pairing_strategy = greedy

With MSA server use, Boltz-2 relies on the MMseqs2 server [85–87] for automatic MSA generation. Output from Boltz-2 used for analyses include predicted structures in CIF format, MSAs, and confidence metrics (confidence score, overall pLDDT, overall pTM, interface pLDDT, interface pTM), as described in the Boltz-2 documentation available on GitHub (https://github.com/jwohlwend/boltz). The Boltz-2 confidence score is calculated as follows: confidence score = 0.8 × overall pLDDT + 0.2 × interface pTM.

For comparison to experimental structures, the RCSB PDB Search API [21] was used to search for structures corresponding to the UniProt IDs of the predicted structures that were deposited after the Boltz-2 model training cutoff date of June 1, 2023 [39]. Search criteria were set to include only structures determined by X-ray crystallography or CryoEM, without mutations, protein only, and containing exactly two chains (corresponding to the two interactor proteins), which retrieved a total of three structures **(Supplementary Figure 1)**. Corresponding AlphaFold3 structures were predicted using the Google DeepMind AlphaFold server (https://alphafoldserver.com/) [28]. DockQ scores and interface RMSD were determined using DockQ v2 [46]. All other molecular analyses, including hydrogen bond and salt bridge determination, hydrophobicity determination, and structure visualization, were performed using ChimeraX [88]. In ChimeraX, hydrogen bonds were determined with the H-bonds tool with default settings (*i.e.,* radius = 0.075 Å and without relaxed distance/angle criteria) and selection of salt bridge only setting allowed for salt bridge determination. Hydrophobic surfaces were determined using the automated molecular lipophilicity potential (MLP) command.

### Pfam domain and interaction network analyses

Pfam domain annotations were curated based on interactor protein UniProt IDs using the InterPro REST API [40]. For a given interactor protein, this included retrieval of Pfam domain IDs and their corresponding start and end positions within the UniProt-obtained protein sequence. To identify Pfam domains less than 5 Å of the interaction interface, the Biopython Bio.PDB package [89] was used to map domain residues within the predicted structural complexes and compute minimum interatomic distances between each domain and the partner interactor protein chain. UniProt biological process keywords corresponding to each interactor protein UniProt ID were obtained using the UniProt website REST API [20]. Chord diagrams of interaction networks were generated using the pyCirclize Python package (https://github.com/moshi4/pyCirclize), with individual proteins plotted as sectors (nodes) along the circumference of the circle and chords (connecting lines) representing interactions with other proteins.

The six biological process networks **(Figure 4e)** were compiled by filtering the entire binary interaction dataset **(Supplementary Table 3)** by UniProt biological process keyword ID, specifically: (1) KW-0053 for apoptosis; (2) KW-0132 for cell division; (3) KW-0221 for differentiation; (4) KW-0234 for DNA repair; (5) KW-0391 for immunity; and (6) KW-0653 for protein transport. The oncogenic network **(Figure 5)** was compiled by filtering the entire binary interaction dataset **(Supplementary Table 3)** by the UniProt disease keyword ID proto-oncogene (KW-0656). Interactions depicted in the chord diagrams in **Figure 4e** and **Figure 5a** are based exclusively on domains < 5 Å of the interaction interface. Genetic variant dbSNP [51] annotations of missense mutations were retrieved using the EMBL-EBI Proteins API [90]. Cross-network analyses were performed by identifying individual proteins that were mutually present in the oncogenic network and in each of the biological process networks separately (*i.e.* oncogenic-apoptosis; oncogenic-cell division; oncogenic-differentiation; oncogenic-DNA repair; oncogenic-immunity; and oncogenic-protein transport). Proteins mutually present within each network pairing served to identify and count interactions involving these shared proteins within the respective biological network, thereby determining the extent of cross-network interaction involvement among the different networks **(Figure 5c)**.

### Statistical Analyses and Graphical Representation

Statistical analyses of medians, quartiles, Spearman correlations, linear regression, as well as corresponding graphical representations shown in the Figures, were performed and produced using GraphPad Prism 10. In Figure 2c-e, *n* = 25 interaction structures which had MSA depths greater than 5,000 sequences are not shown on the graphs, but were not excluded from statistical analyses. The data used for statistical analyses are presented in **Supplementary Table 1** and **Supplementary Table 2**.

### Data Availability

All predicted complex structure models described herein are freely available online at https://github.com/structural-interactome/human-interactome.

## ACKNOWLEDGMENTS

This work was supported by the Levy-Longenbaugh Donor-Advised Fund (to R.P. and W.A.). RCSB Protein Data Bank core operations are jointly funded by the National Science Foundation (DBI-2321666 to S.K.B.), the US Department of Energy (DE-SC0019749 to S.K.B.), and the National Cancer Institute, the National Institute of Allergy and Infectious Diseases, and the National Institute of General Medical Sciences of the National Institutes of Health (R01GM157729 to S.K.B.).

We are grateful to Saro Passaro, Gabriele Corso, Jeremy Wohlwend et al. [39] from the Massachusetts Institute of Technology (MIT) for the development and open-source availability of Boltz-2, which was central to the current work. We would also like to acknowledge the European Bioinformatics Institute (EMBL-EBI) Molecular Networks team for open access to the IntAct database [14] and the UniProt Consortium [20]. Molecular analyses and visualization were performed using UCSF ChimeraX, developed by the Resource for Biocomputing, Visualization, and Informatics at the University of California, San Francisco, with support from the National Institutes of Health R01-GM129325 and the Office of Cyber Infrastructure and Computational Biology, National Institute of Allergy and Infectious Diseases.

## AUTHOR CONTRIBUTIONS

A.M.I., C.M., S.K.B., R.P., and W.A. conceptualization; A.M.I. and C.M. methodology; A.M.I. investigation; A.M.I., C.M., S.K.B., R.P., and W.A. formal analysis; A.M.I. and C.M. writing—original draft; S.K.B., R.P., and W.A. writing—review & editing; S.K.B., R.P., and W.A. funding acquisition; S.K.B., R.P. and W.A. overall project supervision.

