## Supplementary Information for "An AI/ML-modeled structural atlas of the human protein interactome with functional and cancer-focused networks"

**SI contents**

Supplementary Figure 1

Supplementary Figure 2

Supplementary Figure 3

Supplementary Figure 4

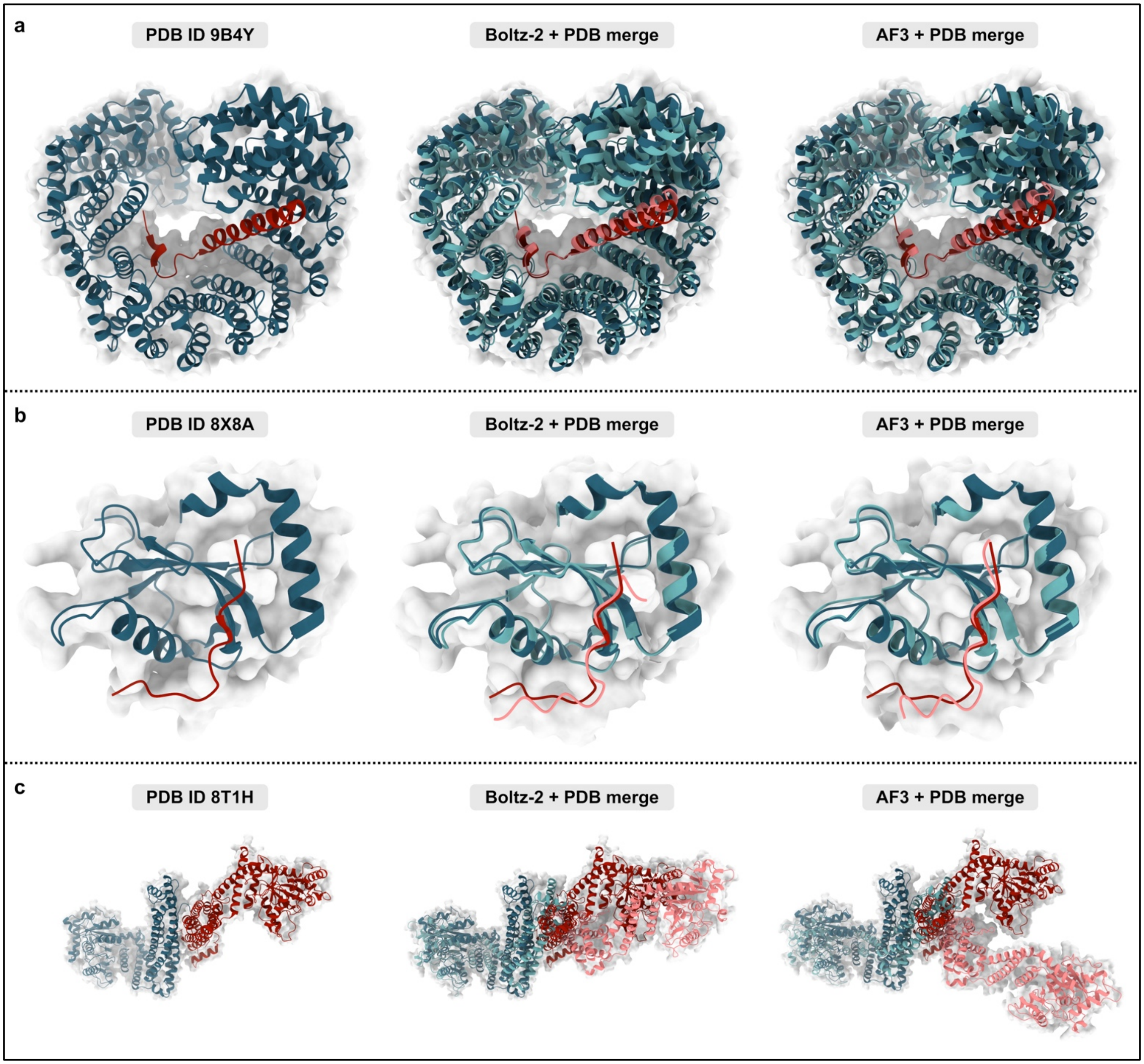


**Supplementary Figure 1. Comparison of predicted structures to experimentally resolved structures not included in Boltz-2 model training. (a-c)** Experimentally determined interaction structures deposited in the PDB beyond the Boltz-2 training cutoff date compared to interaction structures predicted by Boltz-2 and AlphaFold3. Interactor protein A is depicted in teal and interactor protein B is depicted in red, with darker shading for experimentally determined structures and lighter shading for predicted structures. **(a)** CryoEM structure of importin subunit alpha-1 in complex with importin subunit beta-1. Measures of accuracy between the PDB structure and predicted structures were DockQ = 0.755 (interface RMSD = 1.516 Å) for Boltz-2, and DockQ = 0.717 (interface RMSD = 1.811 Å) for AlphaFold3. **(b)** Crystal structure of gamma-aminobutyric acid receptor-associated protein-like 1 in complex with starch-binding domain-containing protein 1. Measures of accuracy between the PDB structure and predicted structures were DockQ = 0.348 (interface RMSD = 5.688 Å) for Boltz-2 and DockQ = 0.372 (interface RMSD = 5.588 Å) for AlphaFold3. **(c)** CryoEM structure of a homodimeric complex of dynamin-1-like protein. Measures of accuracy between the PDB structure and predicted structures were DockQ = 0.210 (interface RMSD

= 3.907 Å) for Boltz-2 and DockQ = 0.201 (interface RMSD = 4.125 Å) for AlphaFold3. In all panels, only residues present in the experimental PDB structures are shown for the corresponding predicted structures.

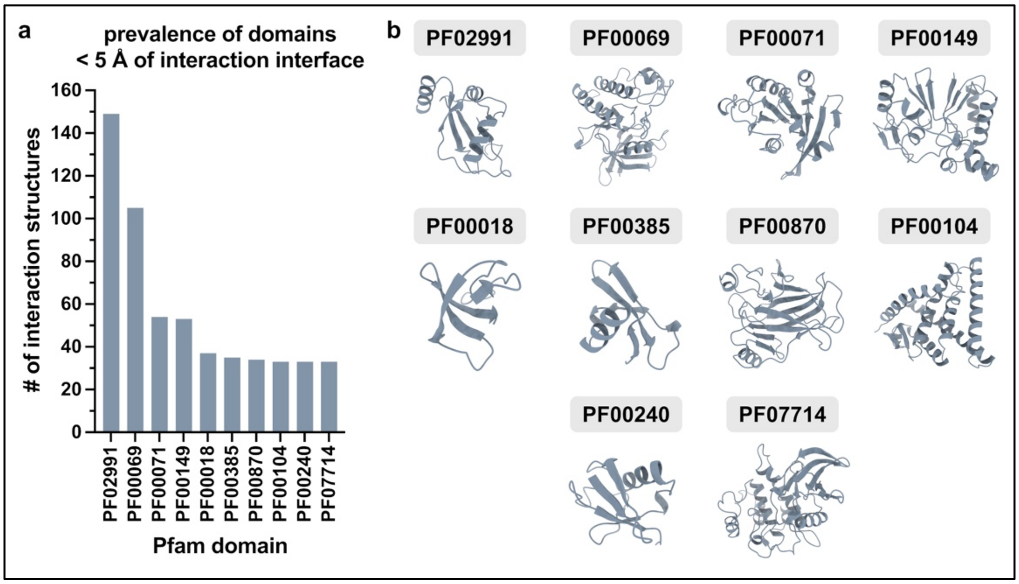


**Supplementary Figure 2. Prevalence of Pfam domains among the predicted interaction structures. (a)** Top 10 most prevalent Pfam domains by number of protein-protein interaction pairs, restricted to only include domains found within 5 Å of the interaction interface. These include: autophagy protein Atg8 ubiquitin like domain (PF02991), present in $n$ = 149 interaction structures; protein kinase domain (PF00069), $n$ = 105 interaction structures; Ras family domain (PF00071), $n$ = 54 interaction structures; calcineurin-like phosphoesterase domain (PF00149), $n$ = 53 interaction structures; SH3 domain (PF00018), $n$ = 37 interaction structures; chromatin organization modifier domain (PF00385), $n$ = 35 interaction structures; P53 DNA-binding domain (PF00870), $n$ = 34 interaction structures; ligand-binding domain of nuclear hormone receptor (PF00104), $n$ = 33 interaction structures; ubiquitin family domain (PF00240), $n$ = 33 interaction structures; protein tyrosine and serine/threonine kinase domain (PF07714), $n$ = 33 interaction structures. The quantification of number of interactor pairs containing a given domain is non-redundant, *i.e.* multiple occurrences of the same domain (either in interactor A or interactor B) do not cumulatively inflate the count. **(b)** Examples of the top 10 most prevalent domains from (a) as found within predicted interaction structures based on Pfam domain sequence annotation.

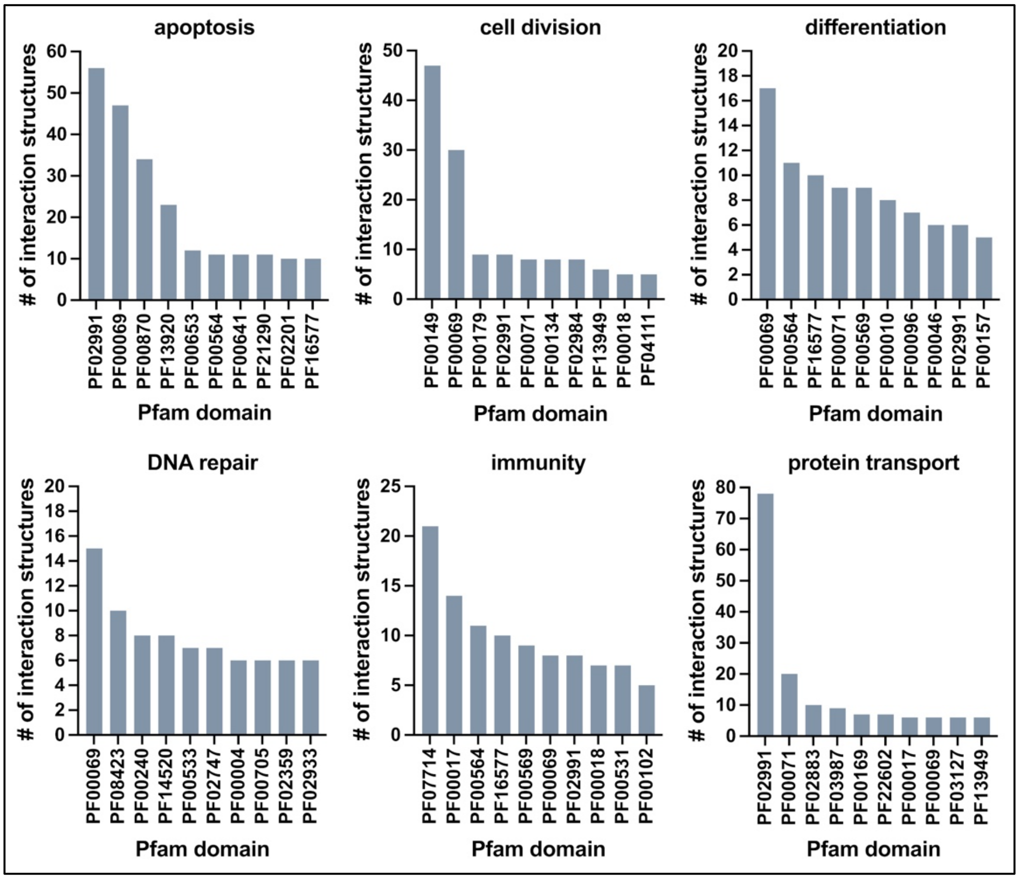


**Supplementary Figure 3. Domain prevalence across the functional interaction networks.** Top 10 most prevalent Pfam domains by number of interaction pairs among the various UniProt biological process keyword groupings. These data were restricted to only include domains found within 5 Å of the protein-protein interaction interface. Group inclusion was based on at least one protein in the interaction pair being annotated with the designated UniProt keyword, including: KW-0053 for apoptosis; KW-0132 for cell division; KW-0221 for differentiation; KW-0234 for DNA repair; KW-0391 for immunity; and KW-0653 for protein transport. The total number of interaction pairs for each group were as follows: apoptosis $n$ = 292; cell division $n$ = 170; differentiation $n$ = 123; DNA repair $n$ = 115; immunity $n$ = 115; and protein transport $n$ = 188.

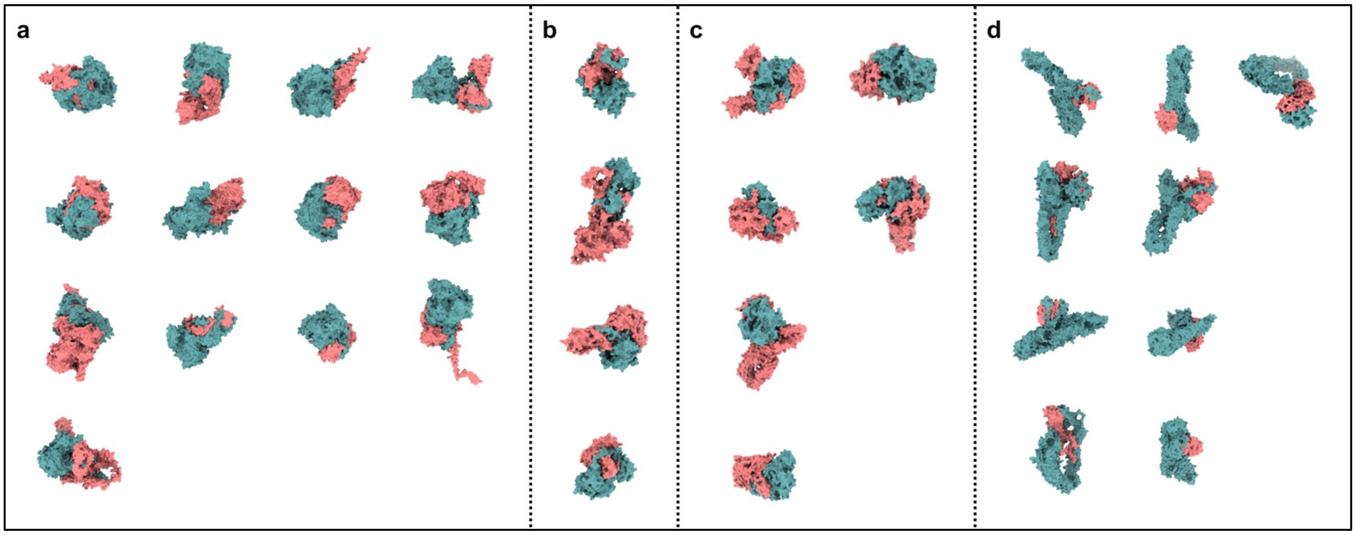


**Supplementary Figure 4. Oncogenic and biological process cross-network protein interaction structures (a-d)** Structures of interactions involving proteins that are mutually present in the oncogenic network and the biological process networks for either differentiation, DNA repair, immunity, and protein transport, corresponding to the cross-network matrix in Figure 5c. These include: **(a)** NAD-dependent protein deacetylase sirtuin-1 (UniProt Q96EB6) within the differentiation network; **(b)** p53 (UniProt P04637) within the DNA repair network; **(c)** Tyrosine-protein kinase Fyn (UniProt P06241) within the immunity network; and **(d)** within the protein transport network. The mutually present (cross-network) proteins are depicted in teal and their interactors are depicted in red.
